# Holistic integration of omics data reveals the drivers that shape the ecology of microbial meat spoilage scenarios

**DOI:** 10.1101/2023.08.31.555692

**Authors:** Simon Poirier, Gwendoline Coeuret, Marie-Christine Champomier-Vergès, Marie-Hélène Desmonts, Dalal Werner, Carole Feurer, Bastien Frémaux, Sandrine Guillou, Ngoc-Du Martin Luong, Olivier Rué, Valentin Loux, Monique Zagorec, Stéphane Chaillou, the ANR Redlosses Consortium group

## Abstract

The use of omics data for monitoring the microbial flow of fresh meat products along a production line and the development of spoilage prediction tools from these data is a promising but challenging task. In this context, we produced a large multivariate dataset (over 600 samples) obtained on the production lines of two similar types of fresh meat products (poultry and raw pork sausages).We describe a full analysis of this dataset in order to decipher how the spoilage microbial ecology of these two similar products may be shaped differently depending on production parameter characteristics.

Our strategy involved a holistic approach to integrate unsupervised and supervised statistical methods on multivariate data (OTU-based microbial diversity; metabolomic data of volatile organic compounds; sensory measurements; growth parameters), and a specific selection of potential uncontrolled (initial microbiota composition) or controlled (packaging type; lactate concentration) drivers.

Our results demonstrate that the initial microbiota, which is shown to be very different between poultry and pork sausages, has a major impact on the spoilage scenarios and on the effect that a downstream parameter such as packaging type has on the overall evolution of the microbial community. Depending on the process, we also show that specific actions on the pork meat (such as deboning and defatting) elicit specific food spoilers such as *Dellaglioa algida,* which becomes dominant during storage. Finally, ecological network reconstruction allowed us to map six different metabolic pathways involved in the production of volatile organic compounds involved in spoilage. We were able connect them to the different bacterial actors and to the influence of packaging type in an overall view. For instance, our results demonstrate a new role of *Vibrionaceae* in isopropanol production, and of *Latilactobacillus fuchuensis* and *Lactococcus piscium* in methanethiol/disylphide production. We also highlight a possible commensal behavior between *Leuconostoc carnosum* and *Latilactobacillus curvatus* around 2,3-butanediol metabolism.

We conclude that our holistic approach combined with large-scale multi-omic data was a powerful strategy to prioritize the role of production parameters, already known in the literature, that shape the evolution and/or the implementation of different meat spoilage scenarios.

## Introduction

Meat products are perishable substrates mainly because of metabolic activities expressed by microbial communities that unavoidably contaminate meat cuts. Once contaminated, meat physicochemical properties, i.e., neutral pH, high water activity and nutrient proficiency, provide excellent conditions for bacterial growth during storage that can result in the production of unwanted colors, textures or odors that contribute to spoilage (Remenant et al., 2015). The microbial ecology of meat products has been the focus of many scientific articles in the last decade (see Nieminem et al., 2012; Chaillou et al., 2015; Hultman et al., 2015; Ferrocino et al., 2016; Fougy et al., 2016; Stellato et al., 2016; Ferrocino and Cocolin, 2017; Wang et al., 2017 as examples). This detailed scrutiny, performed with the use of non-cultural 16S-based amplicon sequencing analyses (also referred to as metataxonomic analyses), led us to revisit the breadth of knowledge covering the diversity of microbiota associated with the spoilage of these perishable foods. In particular, the presence of unexpected species and the detection of some, such as lactic acid bacteria species, usually not or seldom revealed by classical plating methods, gave a new perception of meat microbiota and putative meat spoilers (Chaillou et al., 2015; Saraoui et al., 2016; Andreevskaya et al., 2018; Säde et al., 2020). Much of the microbiological data could gradually be updated and eventually centralized in specialized food databases (de Filippis et al., 2018; Parente et al., 2019), highlighting the predominant role of various psychrotrophic bacterial species belonging mainly to the Proteobacteria (*Photobacterium, Pseudomonas, Psychrobacter, Serratia*) and Firmicutes (*Brochothrix*, *Carnobacterium*, *Lactococcus, Latilactobacillus, Leuconostoc*) phyla, as well as the large variety of assemblies of these bacteria.

However, above all, the integration of all these studies has made it possible to better formulate the key research questions about the microbial ecology that surrounds meat spoilage phenomena. Many biotic parameters (i.e., microbiota present in the raw food, their components and relative abundance) and environmental parameters such as storage conditions (i.e., gas mixtures used for packaging, temperature) and process (i.e., addition of preservative compounds) provide an extraordinary array of functional diversity. Various microbial assemblages between the different groups mentioned above can be observed depending on these parameters. This has led to an overwhelming number of spoilage situations that are hard to tackle and that limit our capacity to produce fundamental knowledge in this research field. A given spoilage microbiota and its functional activity is thus a unique outcome that combines all the variables along a production process, from the time the animal is slaughtered to the final product at the end of its shelf life.

The implementation of large-scale sampling strategies (hundreds of samples) to map the microbial flow along a production process can be very interesting to delineate how microbial communities are structured in a process-dependent manner and the main steps involved in the variations (Stellato et al., 2016; Higgins et al., 2018; Säde et al., 2020; Zagdoun et al., 2020). Moreover, tools for multivariate analysis of a combination of various datasets (Luong et al., 2021) obtained through different approaches such as microbiome, metabolome and different physicochemical or microbiological data may contribute to enriching our knowledge about spoilage occurrence by detecting situations more or less favorable to spoilage development. Once this task completed, it should be easier for food microbiology researchers and process engineers to focus on certain types of microbial communities, to study them in detail, and to look for innovative solutions to reduce their spoilage potential. This strategy can also serve to identify relevant process-specific biomarkers to improve predictive tools for the reliable estimation of the shelf life of fresh meat products. Nevertheless, this type of approach is still not widespread despite its undeniable potential in terms of food safety and meat production sustainability (reducing food waste).

For this purpose, we have produced a large multivariate dataset obtained on the production lines of two similar types of fresh meat products (poultry and raw pork sausages) and have made it available for the scientific community (Poirier et al., 2020). In the present study, we describe a full analysis of this dataset and decipher how the spoilage microbial ecology of these two similar products is nevertheless shaped differently depending on the parameters characteristic of each one.

## Materials and methods

### Dataset presentation

The data presented in this article and their production methods are detailed in our data paper (Poirier et al., 2020). The two meat types were chosen because of their similar recipe (batter of 75-80% ground meat and 15-20% fat mixed with spices, both including potassium lactate as a preservative). Process parameters were subject to controlled variations. The first parameter, the concentration of potassium lactate that is commonly used as a chemical preservative, was modulated in the meat batter: a full normal dose (2% and 1.13% wt/wt, for pork and poultry, respectively; the one generally used in the production facilities that were studied), a half dose, or none. The second parameter, packaging atmosphere, was one of three different types that are also routinely used in the studied factories: air (∼21% O_2_ -∼78% N2), CO_2_-enriched and O_2_-depleted packaging (50% CO_2_ – 50% N2, further referred to as O_2_-packaging), and O_2_/CO_2_-enriched packaging (70% O_2_ – 30% CO_2_, further referred to as O_2_+ packaging).

Several types of analyses were performed for each sample for the purpose of providing a comprehensive microbial ecology of spoilage during storage and to show how the process parameters influence this phenomenon. The analyses include meat type; gas (O_2_ and CO_2_) content in headspace packages, sausage pH, chromametric measurements, plate counts (total mesophilic aerobic flora and lactic acid bacteria), sensory properties of the products, meta-metabolomic quantification of volatile organic compounds (VOCs) and bacterial community metataxonomic analysis. Bacterial diversity was monitored using two types of amplicon sequencing: 16S rDNA V3-V4 and the GyrB encoding gene (Poirier et al., 2019) at different time points during storage for the different conditions (576 samples for *gyrB* and 436 samples for 16S rDNA were obtained). Sequencing data were generated using Illumina MiSeq. The sequencing data were deposited in the bioproject PRJNA522361. Sample accession numbers vary from SAMN10964863 to SAMN10965438 for the *gyrB* amplicon, and from SAMN10970131 to SAMN10970566 for 16S (Poirier et al., 2020).

### Unsupervised statistical analyses

Analyses of alpha bacterial diversity were carried out with phyloseq R package, v 1.36.0 (McMurdie and Holmes, 2013). ANOVA followed by a Tukey test were used to compare alpha diversity and spoilage intensity at different sampling times. Dendrograms that visualized sample clustering according to their bacterial community composition were plotted on the basis of their weighted Unifrac distance (Hamady et al., 2010) and the Ward D2 clustering method (Ward, 1963).

As a global approach, unsupervised principal component analyses (PCA) and correlation circles associated with PCA results were plotted with the ade4 R package, v1.7.19 (Dray and Dufour, 2007). Centered scaled PCA results were performed with sensory analysis and volatile organic compound quantification datasets. Bacterial community compositions within meat samples were first ordered according to the Bray-Curtis dissimilarity index values using a principal coordinate analyses (PCoA) and then plotted using the ggplot2 R package, v 3.3.6 (Wickham, 2016).

### Supervised statistical analyses

The MixOmics R package, v 6.17.29 (Rohart et al., 2017), was used to implement supervised partial least square discriminant analysis (PLS-DA) and plot sample distribution according to their bacterial population composition and to preliminary defined groups on factorial planes. The objective of this supervised approach was to more precisely target robust bacterial indicators associated with variations in environmental parameters. For this analysis, data were transformed with centered log ratio (CLR) transformation to account for compositional structure of the scaled data (Aitchison, 1982). Briefly, the log (base 2) of each count divided by its corresponding geometric mean was calculated. Performance of the PLS-DA model was checked using 5-fold cross validation repeated 10 times (*perf* function). This analysis enables the calculation of an error rate that targets the number of components to be kept. The number of components corresponding to the best error rate was then selected. A sparse PLS-DA (*sPLS-DA* function) enabling the selection of the most discriminant bacterial populations (variables) within PLS-DA results, was also conducted with this package. This selection is also based on the calculation of an error rate according to an evaluation of the PLS-DA model (*tune.splsda* function) using 5-fold cross validation repeated 10 times. The number of variables corresponding to the best error rate is then selected.

### Correlation analyses between heterogeneous multi-omics datasets

Dataset integration was performed using Data Integration Analysis for Biomarker discovery, DIABLO (Singh et al., 2019), implemented in the mixOmics framework. In our correlation analysis, we combined a centered log ratio (CLR) normalized relative bacterial abundance table aggregated at the species level using the *tax_glom* function of the phyloseq R package with the CLR normalized volatile organic compounds (VOCs) abundance table. DIABLO extends sparse Generalized Canonical Correlation Analysis (sGGCA) for multi-omics and supervised integration. It performs multivariate dimensionality reduction and selects correlated variables (based on latent component scores) across datasets. Feature selection is performed internally using lasso penalties. The data are then projected into a smaller dimensional subspace spanned by the components for prediction. Tuning of the component and variable number were performed using 5-fold cross validation repeated 10 times. The number of variables corresponding to the best error rate is then selected.

### Construction of the ecological network

Construction of the ecological network was performed with Gephi software, v 0.9.2 (Bastian et al., 2009). The node and edge table was configured from the subset mixDIABLO analysis (see Results section) using hit frequency and cumulative r values to weigh the edges. The network was constructed using the forceAtlas2 algorithm (Jacomy et al., 2014).

## Results

### Summary overview of the dataset and of the analysis strategy

Technical details about the experimental design can be found in our data article (Poirier et al., 2020). Briefly, we monitored the production of two meat products: poultry and pork chipolata-type sausages. We assessed how uncontrolled input characteristics (meat type) and associated microbiota influence the spoilage of raw sausages, while two additional process variables (potassium lactate concentration and packaging atmosphere) were subjected to controlled variations (see Material & Methods section).

The large-scale sampling involved ten batch replications spanning six months of production and made it possible to gather over 360 samples per meat type along the production chain (10 replicates x 3 packaging conditions x 3 lactate concentrations x 4 sampling times). Sausages were stored until day 22, mimicking the conditions used for use-by-date determination, and sampled at day 2, 7, 15 and 22. On the basis of these samples, we produced multivariate data from the monitoring of the physicochemical parameters (pH of sausage, gas mixtures in the pack head space, sausage color determined by chromametry), bacterial loads, microbiota diversity, including both 16S (taxonomic assignment of operational taxonomic units at the genus level (Poirier et al., 2018)) and *gyrB* (taxonomic assignment of operational taxonomic units at the species and subspecies level) amplicon sequencing, metabolomic analysis of VOCs, and sensory analysis using six descriptors evaluated by a trained panel.

### Bacterial growth and alpha diversity dynamics depend on the raw food input (meat type)

The analysis of microbial α-diversity showed that the total number of OTUs initially contaminating pork (Figure 1A) and poultry sausage (Figure 1B) were relatively similar (196 ± 43 and 185 ± 25 OTUs, respectively). However, the initial contamination level of poultry sausage (5.8 ± 0.6 log_10_ CFU/g) was significantly higher than that of the pork samples (4.3 ± 0.7 log_10_ CFU/g) (Poirier et al., 2020). This higher contamination level found in poultry was also associated with a faster bacterial growth dynamics during the storage of poultry sausages (Figure 1D). Indeed, the population level enumerated in poultry sausages reached 8.6 ± 0.6 log_10_ after 7 days and then stabilized around these values, whereas in pork sausages, the population level reached 8.3 ± 0.5 log_10_ after 15 days (Figure 1C) and stabilized only after 22 days (Poirier et al., 2020).

**Figure 1.**
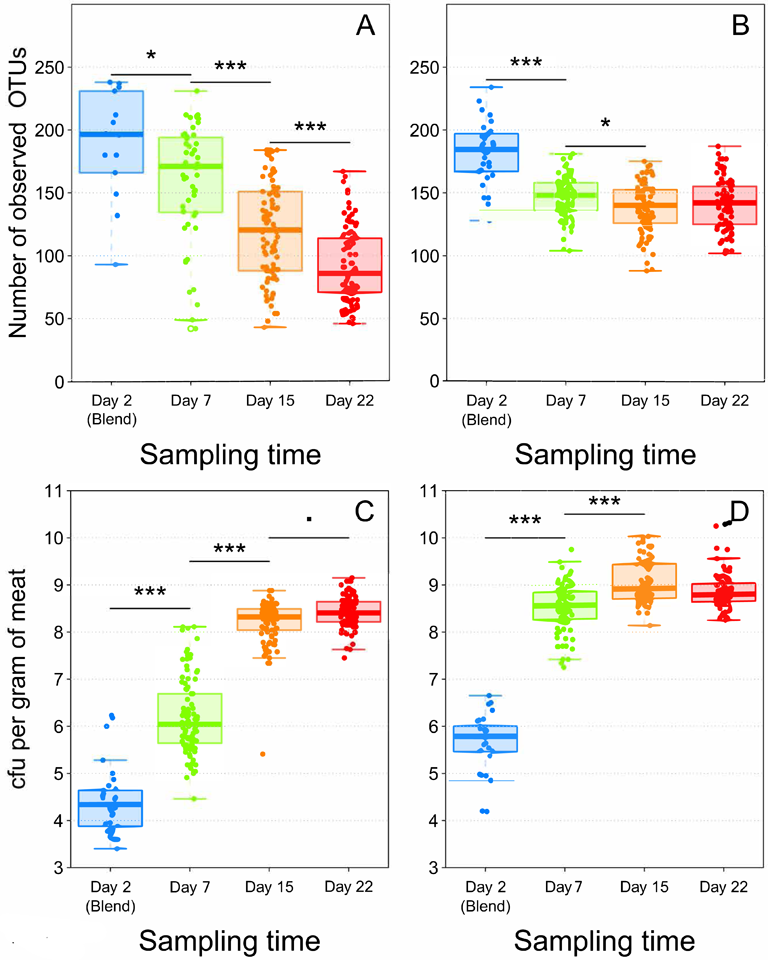
Boxplot analysis of both *gyrB* alpha diversity (number of observed OTUs) (Panels A and B) and of bacterial growth in log_10_ CFU/g (Panels C and D) according to sampling time. Panels A and C = Pork samples. Panels B and D = Poultry samples. Number of samples analyzed = 538 for both food matrices (240 for pork and 298 for poultry). Statistical analysis performed with ANOVA followed by a Tukey test (*** for *p* < 0.001; * for *p* < 0.01; · for *p* < 0.05).

The evolution of microbial α-diversity was consistent with bacterial growth, similarly showing 7 days of difference in dynamics between the two meat types. Within poultry sausages, diversity decreased down to 148 ± 16 OTUs after 7 days, and stabilized around these values until the end of the storage experiment at 22 days. On the other hand, a greater simplification of α-diversity in pork sausages produced a linear decrease during storage down to 86 ± 30 OTUs after 22 days (∼ 40% lower than for poultry).

These results clearly illustrate the selective diversity process related to the exponential growth phase of the bacterial community, a process that stops when this community reaches the stationary phase. The differences observed between the two sausage types may result from differences in the nature of meat: animal type (feathered bird vs. mammal), slaughtering, evisceration, butchering, and cutting practices are different for poultry and pork and may differently influence initial contamination.

### Intrinsic food type characteristics also drive the bacterial community structure

Beyond the number of bacterial cells and species present in the sausage samples, we were also interested in identifying the species making up these communities. As shown in Figure 2A, hierarchical clustering based on weighted Unifrac distance, revealed that, bacterial communities found in pork sausages and poultry sausages greatly differed in their structure, except in a few of the samples. However, *Latilactobacillus curvatus* and *Leuconostoc carnosum* were two species constantly recovered among the dominant bacteria in both food products and might be considered as a bacterial signature of this type of meat product. Bacterial community structure of poultry sausages confirmed the results shown in Figure 1, indicating much higher species diversity than that of pork sausages. Poultry sausage samples also showed considerable inter-sample variability with at least three main branches in the compositional clustering tree.

**Figure 2.**
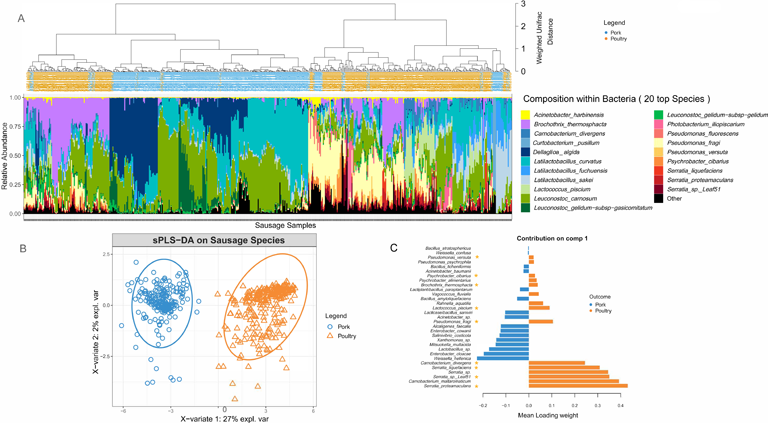
Clustering of sausage samples and determination of sausage-type specific populations. **Panel A.** Clustering of sausage samples according to the composition of the bacterial community. Analysis was only performed with samples collected at 8, 15 and 22 days (blend samples and sausages at day 2 were discarded because of the small number of pork samples with enough DNA and of the high number of species detected in early samples but that were no longer found later on). Before analysis, diversity data were merged at the species level. Overall, the dataset was composed of 494 samples encompassing 149 species. Clustering was performed with weighted Unifrac distance and Ward’s clustering algorithm. Relative abundance of the TOP 20 species is shown with a color code (blue/green gradient for Firmicutes and red/yellow gradient for Proteobacteria). **Panels B and C**. Sparse partial least square discriminant analysis (sPLS-DA, see Materials & Methods) on the microbial diversity of sausage samples with discrimination based on meat type. The plot in **panel B** shows the distribution of samples along the two principal components (X-variate 1 and 2) and the weight of each on the overall variation observed. The plot in **Panel C** shows all the bacterial species with a significant loading weight on the distribution of samples along component 1. Asterisks indicate the poultry vs. pork discriminating species that belong to the dominant population.

A sparse partial least square discriminant analysis (sPLS-DA) was performed to statistically confirm the differences observed in Figure 2A. It enabled the selection of the most discriminative species between the two types of meat. Spatial distribution of sausage samples on a factorial plane based on the relative abundance of these selected species is illustrated in Figure 2B. This representation showed that both meat types were exclusively separated along the first component, indicating that meat type is the most important variable explaining the clustering of samples based on bacterial community structure. A total of 31 species were identified as being able to discriminate the two sausage types. Figure 2C shows a classification of these species according to their discriminating weight. The discriminating species in pork sausage all belonged to the sub-dominant bacterial population (not visible among the top 20 species shown in Figure 2A), indicating that the dominant bacterial population of pork sausages is also found in poultry sausages.

Conversely, many of the discriminating species in poultry sausages were identified as being part of the dominant population. Furthermore, these dominant poultry populations showed considerable relative abundance variations among samples, suggesting that intrinsic meat characteristics structured these communities. These results prompted us to separate the pork dataset from that of the poultry dataset in order to further assess the influence on bacterial community structure and dynamics of the variables we chose: potassium lactate concentration and packaging atmosphere during storage.

### Packaging atmosphere has a strong structuring effect on the bacterial community of poultry sausages

Weighted Unifrac hierarchical clustering performed on a poultry sample subset (Figure 3A) revealed that the three main branches of the tree separating the various bacterial community structures were strongly associated with the three different packaging atmospheres. Statistical sPLS-DA analysis (Figure 3B) confirmed this trend and made it possible to discriminate samples according to the bacterial species that were the most affected by the packaging atmosphere. Our analysis revealed that samples initially stored under air packaging were discriminated along the first component. This discrimination was exclusively explained by the strong domination of aerobic *Pseudomonas* species, in particular, the psychrotrophic species *Pseudomonas fragi* and *Pseudomonas psychrophila.* sPLS-DA confirmed that discriminant populations found in samples stored under O_2_+ conditions were mainly assigned to *Brochothrix thermosphacta, Psychrobacter* and *Leuconostoc* species, whereas *Lactococcus piscium* and *Hafnia alvei* were discriminating species of anaerobic storage conditions (Figure 3C). We noted that cluster separation on the sPLS-DA factorial plane was less marked for air-stored samples than for the two other types. This phenomenon was notably linked to the fact that under air packaging, a progressive shift to anaerobic conditions already occurred at 7 days of storage because of oxygen consumption associated with CO_2_ production by the microbiota (data available in our data paper (18)). Overall, our results demonstrate that although there is a fast microbial growth and early stabilization of the microbial population level in poultry sausages (∼7 days), the packaging used for storage mediates strong dynamic structuring.

**Figure 3.**
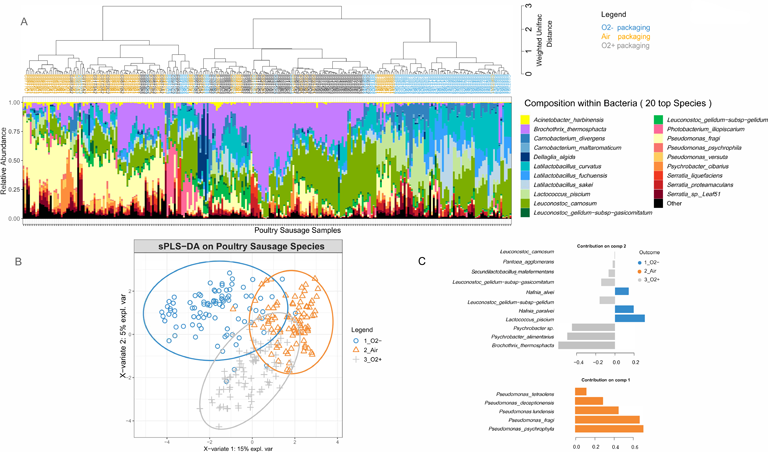
Clustering of poultry sausage samples according to the species composition of the bacterial community in correlation with atmosphere packaging. **Panel A.** Clustering was performed with weighted Unifrac distance and Ward’s clustering algorithm. Samples are colored according to the packaging type. **Panels B and C**. Sparse partial least square discriminant analysis on the microbial diversity of poultry sausage samples with discrimination based on packaging type. The plot in Panel B shows the distribution of samples along the two principal components (X-variate 1 and 2) and the weight of each on the overall variation observed. The plot in Panel C shows all the bacterial species with a significant loading weight on the distribution of samples along component 1 and component 2.

Finally, we performed three sub-settings of the poultry sausage dataset according to the packaging parameter. These sub-settings aimed at evaluating the role of the lactate concentration (third parameter) on additional structuring of the bacterial community in poultry sausages. However, this strategy could not reveal any effect of lactate concentration (analysis not shown). A similar analysis performed on pork samples did not reveal any similar structuring effect of atmosphere packaging on bacterial populations (data not shown).

### The dynamics of bacterial communities in pork sausages are strongly dependent on how the meat is pre-processed before sausage production

Surprisingly, the bacterial dynamics in pork sausage samples evolved differently than those of poultry sausage samples. Indeed, although packaging did not reveal any bacterial community structuring effect, we observed a strong bimodal clustering of the meat batches over time. As shown in Figure 4 upper panels, two groups of batches (P1, P3, P4, P5 vs. P2, P6, P8, P9, P10, and P11) could be distinguished. Differences in terms of community structure increased over storage time, independently of the packaging or lactate parameter. Thorough scrutiny of the metadata associated with the sampling strategy revealed that this binary structuring was correlated with the way the pork meat was pre-processed before being prepared for the meat blend (Figure 4 lower panels).

**Figure 4.**
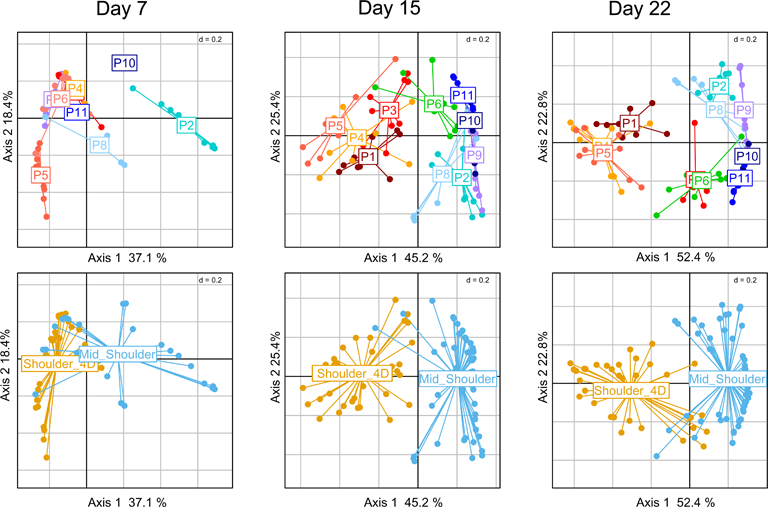
Clustering of pork sausages in correlation with storage time and primary cuts used for sausage making. Principal coordinate analysis (PcoA) of pork sausage samples using the Bray-Curtis dissimilarity index. From left to right, several PcoA analyses are shown on the sub-dataset representing the storage time. On upper panel, the samples are colored according to the ten meat batches analyzed (P1-P11, each dot representing a sausage sample, and on lower panels, the samples are colored according to the type of primary meat cuts used for sausage making.

In general, mid-shoulder meat is used to prepare pork sausages, but in some cases, additional deboning and defatting processes are applied in order to improve meat quality. This type of meat is usually referred to as shoulder 4D (boneless without rind or shank and fat trimmed). As shown in Figure 5, the main consequence of this pre-process step leads to the dominance of the bacterial species *Dellaglioa algida* (formerly referred to as *Lactobacillus algidus*), a well-known psychrothrophic meat-borne spoilage lactic acid bacterium (Jääskeläinen et al., 2016; Andreevskaya et al., 2018; Mansur et al., 2019; Säde et al., 2020). This may result from an initial contamination by this species as a consequence of additional meat processing steps, whereas mid-shoulder samples are dominated by *Leuconostoc carnosum* and *Latilactobacillus curvatus*. Further breakdown of the pork sausage samples according to the type of shoulder did not reveal any significant influence of packaging or lactate concentration on the bacterial community structure (data not shown).

**Figure 5.**
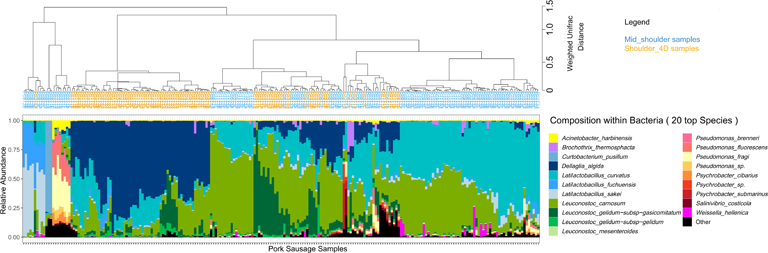
Clustering of pork sausage samples according to the species composition of the bacterial community. Samples are colored according to the primary meat cuts. Clustering was performed with weighted Unifrac distance and Ward’s clustering algorithm.

### Sensory spoilage detection is time- and packaging-dependent.555243v1

Next, our analytical strategy focused on the spoilage characteristics of the sausage samples in order to investigate the role of the different parameters on this phenomenon and to estimate the possible correlations of the various structures of the bacterial communities with spoilage. The kinetics of the global spoilage intensity for pork and poultry sausages during storage are shown in Figure 6A and Figure 6B, respectively. These boxplots indicate a clear correlation between storage time and spoilage sensory intensity for both products. Furthermore, spoilage intensity of poultry sausages after 7 days of storage was comparable to the level reached by pork sausages after 15 days of storage. This faster spoilage kinetics thus corroborates the higher contamination level and the faster microbial growth during storage observed in poultry sausages. Nevertheless, we next investigated in more detail the various spoilage sensory descriptors evaluated by the sensory panel (ethereal, sulfurous, prickly, rancid, old cheese or fermented).

**Figure 6.**
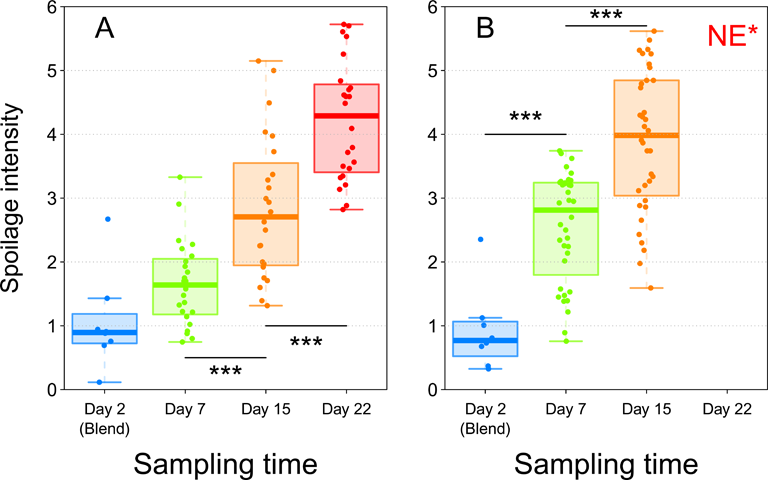
Boxplot analysis of sensory data for both pork (Panel A) and poultry (Panel B) sausages according to storage time. Trained members of the sensory jury gave intensity values (from 0 to 6) to describe the global spoilage intensity. For every sample, the values of this descriptor were averaged between the juries. In total, 160 samples (80 for pork and 80 for poultry) were analyzed with the seven descriptors. Statistical analysis was performed with ANOVA followed by a Tukey test (*** for *p* < 0.001). *NE: Not Evaluated, because these samples were all strongly spoiled and not suitable for sniffing analysis.

These descriptors were analyzed for both pork and poultry sausages together and the results are presented in Figure 7. Among the various parameters tested, neither the meat type nor the packaging atmosphere seemed to significantly influence the sensory spoilage intensity (Figure 7). On the other hand, all sausage samples were discriminated along the factorial plane according to the storage time (60% of variance) i.e., spoilage increases with time, but also depends on other variables. In fact, we noticed that sausage samples located on the right part of the PCA factorial plane showed little discrimination between them. Indeed, these early storage time samples were considered as being non-spoiled (little sensory variation). On the other hand, late storage time samples showed more obvious discrimination along the second axis (12.8% of variance). No influence of lactate concentration was observed (data not shown).

**Figure 7.**
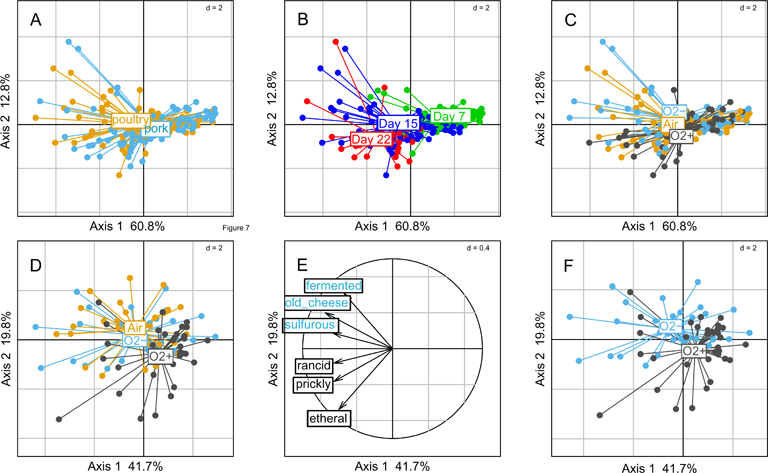
Principal component analysis of sausage spoilage based on six specific quantitative sensory descriptors (1 – ethereal, 2 - sulfurous, 3 - prickly, 4 - rancid, 5 - old-cheese, 6 - fermented odors given by the jury as described in the caption of Figure 6 for global spoilage intensity) and after reduced centered normal distribution (normalization). On the plots of Panels A, B and C, samples are colored according to meat type, storage time and gas mixture used for packaging, respectively.

### The volatile spoilage metabolic profiles of meat sausages corroborate the sensory analyses

Our sensory analysis results prompted us to investigate in more detail the metabolic profiles of VOCs found in the food samples and resulting from bacterial metabolism. Overall, total VOC quantification by gas chromatography coupled with mass spectrometry (GC-MS) enabled the detection of 22 molecules (Figure 8). Our first analysis compared the quantification of these molecules, from the control (fresh sausage) to those of samples at the end of storage (22 days) when molecules are assumed to have reached higher concentrations. For seven molecules (sabinene, alpha-pinene, acetone, alpha-terpinene, alpha-phellandrene, alpha-thujene and hexanal), the concentration was not significantly different at these two times. Thus, these VOCs were estimated to be natural molecules present in the sausage ingredients (meat or spices) and not produced by the microbiota. For the remaining 15 molecules, we observed a significant production during storage, and differences could be detected between pork and poultry sausages.

**Figure 8.**
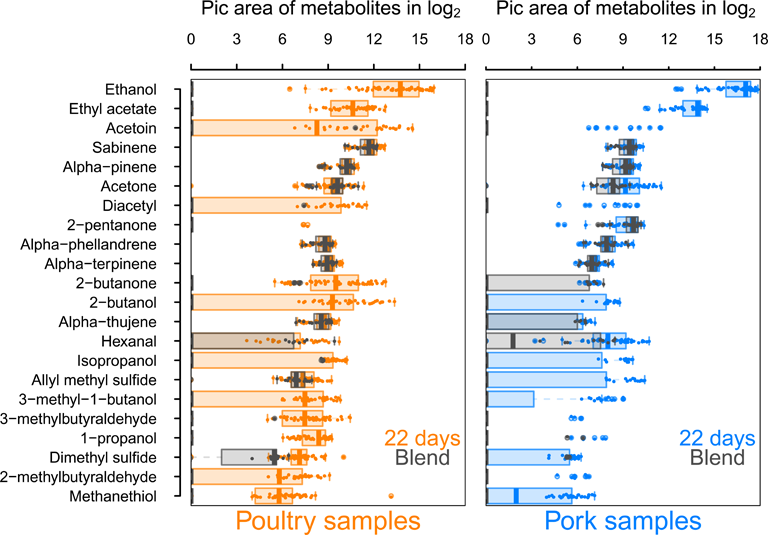
Comparative analysis between blend samples and sausage samples at 22 days of storage of 18 VOCs significantly detected in both types of samples. Results for poultry sausages are colored in orange (left panel) and those for pork sausages are colored in blue (right panel). Sausage samples (day 2) are shaded in gray and printed on the top of samples analyzed after 22 days of storage to compare whether the production during storage is significant and to discriminate compounds naturally present in meat, independently from bacterial growth and metabolism. Molecule quantity is shown in log2 of pic area and VOCs are listed, ranging from the most concentrated (ethanol) to the least concentrated (methanethiol).

Among these differences, we noticed that some molecules such as 2-pentanone and 2-butanone are already present in large amounts in fresh pork sausages but not in fresh poultry sausages. The reverse situation occurs for allyl methyl sulfide. The most abundant molecules were ethanol and ethyl acetate, with an average production level about 1.8-fold higher in pork sausage than in poultry sausage. In contrast, the diversity and production level of all other molecules were higher in poultry sausages. Thus, VOC characterization clearly revealed the differentiated microbial metabolic activities between both types of meat.

Correlations between the 15 discriminant VOCs and the packaging atmosphere were also identified by PCA at each sampling time and for both meat products (Figure 9). Interestingly, the packaging influence on VOC production begins earlier (7 days) during storage time for poultry sausages than for pork sausages (22 days). This corroborates our previous observation revealing that packaging has a more direct influence on the microbiota of poultry sausages than that of pork sausages. The PCA also revealed that, as already demonstrated for the bacterial diversity analysis and sensory analysis, in samples stored with air positioned between the two other groups, it is the result of the progressive switch from aerobiosis to anaerobiosis. Nevertheless, VOC analysis revealed a stronger clustering depending on the packaging condition than did sensory analysis. Lactate concentration did not impact VOC production (data not shown).

**Figure 9.**
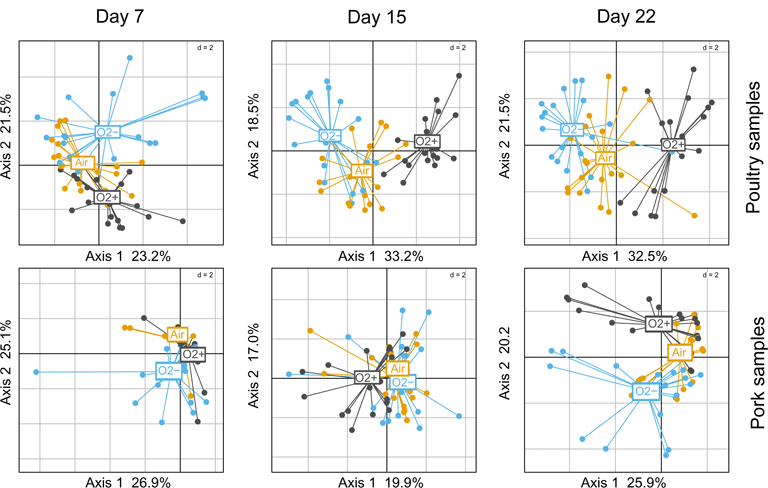
PCA analysis using reduced centered normal distribution (normalization) of VOC profiles (using only the 15 discriminating compounds) between sausage samples. Poultry samples are on the top; pork samples are at the bottom. Three PCAs were carried out from 7 to 22 days of storage. Samples are colored according to packaging.

Correlation circles presented in Supplementary Figure 1 were plotted with the most discriminated samples stored under O_2_+ or O_2_-packaging and collected at different storage times. They enabled the identification of the specific influence of each VOC on the spatial distribution of samples on the factorial plane. Notably, they highlighted that for both types of meat products, compounds such as methanethiol, dimethyl sulfide and allyl methyl sulfide were specifically associated with O_2_-packaging. This confirms the high “sulfurous” ranking attributed by the sensory panel to these samples. Moreover, 1-propanol, 2-butanol and 2-butatone were also correlated with O_2_-packaging for both meat products.

Together with ethanol and ethyl acetate (produced under all conditions but at higher levels under O_2_-packaging), they might also contribute to the fermented alcoholic sensory description associated with this packaging. VOCs related to O_2_+ packaging were also consistent for both types of sausages. Acetoin, diacetyl and 3-methyl-1-butanol were strongly associated with this packaging, thus corroborating the “rancid” and “ethereal” sensory description of these samples. Finally, compounds such as isopropanol, 2- and 3-methylbutyraldehyde had a more complex pattern of production and could not be significantly associated with either packaging, storage time or type of meat.

### Metabolic ecological network reconstruction reveals that bacterial communities are structured on the use of six main spoilage catabolic pathways

The results described above suggested a strong correlation between VOC concentration, bacterial community structures and packaging. To further investigate the role of each bacterial community in the metabolic processes leading to the production of these molecules, we carried out the reconstruction of an ecological network on the basis of a matrix of correlations between bacterial species abundance and VOC concentrations.

We performed several mixDIABLO analyses from the MixOmics package, a tool dedicated to highlighting the correlation between heterogeneous multi-omics datasets, especially when a high number of repeated measurements are available. Overall, we were able to collect 317 sausage samples for which both bacterial species abundance and VOC concentrations were dynamically collected during storage. However, since each variable of the experimental design (storage time, meat type, lactate concentration and packaging type) could represent a major source of variation, we supervised several variable-driven subset analyses to guide the integration process. For each subset, positive correlation values from (*r* ≥ 0.5) were collected to produce a matrix of correlations between the two types of omics data (bacterial species and VOC abundances; see design in Supplementary Figure S2). On the basis of the analysis of 36 subsets, only 21 enabled the identification of 233 significant positive correlations involving 27 bacterial species and 14 VOCs. As shown in Supplementary Figure S2, comparative analysis between O_2_- and O_2_+ packaging revealed the highest number of correlations, whereas the comparative analysis of air and O_2_+ packaging failed, in most cases, to produce such correlations, except for the end of storage at 22 days when air packaging has probably switched to anaerobiosis. Similarly, data from the end of storage (22 days) produced more significant correlations than those from the beginning of storage (7 days) since metabolites accumulate over time and microbial diversity decreases. Finally, data from poultry sausages provided more correlation hits than those from pork.

However, combining both datasets allowed the breakthrough of about 30% new correlations not seen in the individual meat type subsets. Redundant correlations between the various mixDIABLO subset analyses were merged to produce 113 unique correlations. The sum of *r* values corresponding to these redundant correlations were used to assign a weight for each of these 113 unique correlations. Using these data, we built an ecological network of interactions (Figure 10). We then cross-referenced the correlations obtained by this network with the genomic knowledge of the identified species and the metabolic pathways listed either in the literature or in specific databases (i.e., KEGG or MetaCyc) in order to identify the main metabolic activities. The network revealed six main independent metabolic pathways involved in the synthesis of VOCs associated with the sensory profile of spoiled sausages with specific associations of bacterial species for each of them. Globally, Firmicutes were the largest group associated with spoilage, with mainly lactic acid bacteria under O_2_-conditions and *B. thermosphacta* under O_2_+ packaging. Only *Pseudomonas* and *Psychrobacter* were correlated with spoilage under O_2_+ packaging.

**Figure 10.**
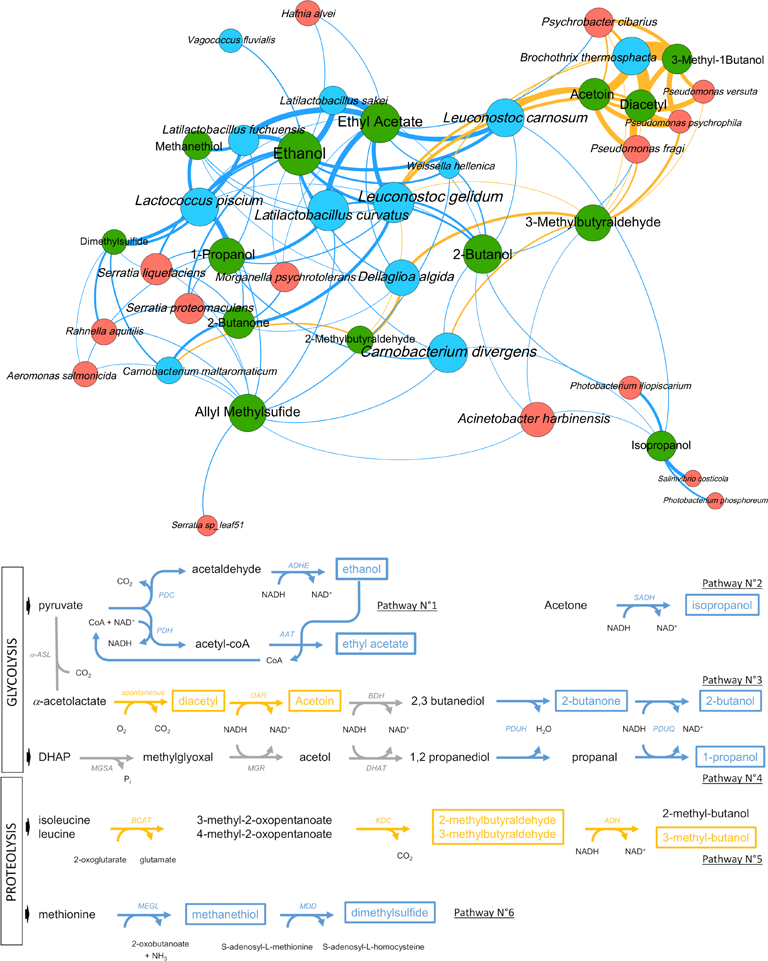
Ecological network showing positive correlations between VOC concentration and microbial taxa abundance and proposed main pathways causing raw sausage meat spoilage. Upper part. Metabolites are shown as green circles and microbial taxa as either blue circles (Firmicutes) or orange circles (Proteobacteria). Links depict the positive correlations based on multiple sPLS-DA mixDIABLO analysis (see main text and Methods section). The thickness of the link is proportional to the strength of the correlation (frequency and *r values*) and the color highlights the packaging modality (blue for O_2_- and yellow for O_2_+). The size of each node (circle) is proportional to the number of correlations. **Lower part.** Outline of the six main pathways producing the VOCs associated with spoilage (a subset of ecological networks is available in Supplementary Figure S3 for each pathway). As above, colors are according to the packaging modality. Detected VOCs are shown with a frame, whereas gray (arrows) or black (VOCs) colors such as the one used for 2,3 butanediol highlight the intermediate position of this compound between two sub-pathways, or compounds that are not detected. Code for enzymatic reactions are shown for every catabolic step: (PDC) pyruvate decarboxylase; (PDH) pyruvate dehydrogenase;(ADHE) bifunctional acetaldehyde/alcohol dehydrogenase; (AAT) alcohol acetyltransferase; (α-ASL) α-acetolactate synthase; (DAR) diacetyl reductase; (BDH) butanediol dehydrogenase; (PDUH) propanediol dehydratase; (PDUQ) propanal reductase; (MGSA) methylglyoxal synthase; (MGR) methylglyoxal reductase; (DHAT) 1,2 propanediol dehydrogenase; (BCAT) α-ketoglutarate-dependent branched-chain aminotransferase; (KDC) 2-keto acid decarboxylase; (ADH) alcohol dehydrogenase; (MEGL) methionine-α-lyase; (MDD) methanethiol methyltransferase.

First of all, the pathway related to ethanol and ethyl acetate production was strongly correlated with lactic acid bacteria, in particular, *Latilactobacillus* and *Leuconostoc* species, with an increased rate under the O_2_-modality.

The second pathway involves the anaerobic production of isopropanol by sub-dominant *Vibrionaceae,* probably from the acetone that is naturally present in large quantities in both types of sausages (see Figure 8). The third pathway connects two sub-pathways and possibly the role of two different microbial communities acting successively. It comprises the aerobic production of diacetyl and acetoin by *B. thermosphacta, Pseudomonas*, and *Leuconostoc* species. In principle, this sub-pathway ends up in the production of 2,3-butanediol, although this VOC was not detected in the sausage samples. However, 2,3-butanediol may also be the substrate for 2-butanone and 2-butanol production, which, according to our results, was strongly correlated with O_2_-packaging and with *L. curvatus* and *Leuconostoc gelidum*. The succession of these two pathways may be particularly efficient under air packaging, where a switch progressively occurs through O_2_ consumption to CO_2_ production. The fourth pathway corresponds to the anaerobic production of 1-propanol from 1,2-propanediol involving mainly *L. curvatus* and *L. piscium*. Like 2,3-butanediol described above, 1,2-propanediol, usually produced from acetol and the methylglyoxal reductase pathway in lactic acid bacteria, was not detected in the spoiled samples, indicating a rapid conversion of this molecule to 1-propanol.

The two remaining pathways involve catabolism of amino acids. It comprises the conversion of leucine/isoleucine to 2- or 3-methybutaraldehyde and 3-methylbutanol under aerobiosis. Correlation analysis revealed that *B. thermosphacta, Pseudomonas* and *Leuconostoc* species are the main producers of these compounds, whereas *D. algida* and *Carnobacterium* species may also contribute. Finally, anaerobic catabolism of methionine to methanethiol and dimethyl sulfide was strongly correlated with *L. piscium*, *Latilactobacillus fuchuensis* and, to a lesser extent, to sub-dominant *Enterobacteriales.* On the other hand, the production of allyl methyl sulfide, a sulfurous compound linked to garlic (used as a spice ingredient in the sausages), was not associated with any known microbial pathway. The positive correlations with many of the bacterial species identified may suggest an indirect effect of bacterial metabolism on the release of this compound from the spice mix.

## Discussion

Meat spoilage is a phenomenon driven by an intricate combination of biotic, environmental, and process parameters. To decipher the spoilage routes leading to various spoilage situations, we produced a large multivariate dataset obtained on the production lines of two similar types of fresh meat products (poultry and raw pork sausages) and by working together with two production facilities in France. Our objective was to prioritize the parameters, already known in the literature, that shape the evolution and/or the implementation of different spoilage scenarios.

Our study highlighted the robustness of the primary bacterial community that was initially structured on the fresh meat batter and by the pre-process implemented to prepare raw meat. Our data demonstrate, despite the variations detected in each of the ten batches monitored during the six months of sampling, that the bacterial communities in poultry meat are distinct from those in pork. Packaging atmosphere (the most downstream parameter in the production) could affect the microbial assemblage and activity of poultry sausages due to the major presence of Proteobacteria, a group encompassing species sensitive to oxygen and CO_2_ levels. On the other hand, in pork sausages that were mainly contaminated by lactic acid bacteria, community structuration remained unchanged during storage and according to packaging type, whereas variations could be observed in their metabolic activity. Interestingly, in both products, modulations of lactate concentration used as a preservative did not influence either the structuration or the metabolic activity of bacterial communities.

The use of metataxonomic data has often been put forward by the food microbiology community as a promising approach to better understand microbial flows along the food chain (Ferrocino and Cocolin, 2017; De Filippis et al., 2018). The development of predictive tools for spoilage or tools for monitoring a production line is often considered as a major stake inherent in this type of approach. However, our results show that this challenge can only be met with an ambitious sampling strategy that makes it possible to elucidate not only the impact of the variables associated with the different production routes, but also to delineate the chain reaction associated with each variable (meat > process > packaging). Moreover, our data demonstrate, on the one hand, that the initial microbial community composition may or may not directly influence the effect provoked by the most downstream parameter (packaging), and that some process parameters (e.g., deboning and defatting of pork shoulder) may elicit specific contamination by food bacterial spoilers (*D. algida*). For this reason, and despite similarities between the two meat products that were analyzed in this study, we can conclude that the construction of general predictive tools for spoilage of meat products remains a difficult objective to reach. From our point of view, each production unit in the world may harbor a characteristic microbial ecology associated with each of its associated processes.

To strengthen our point of view, we have recently proposed another exploratory statistical workflow on the poultry data subset described in this article (Luong et al., 2021). This approach, which used multiblock path modeling workflow (Cariou et al., 2019), already showed causal links between some bacterial species and some VOCs. The integrated holistic approach developed here appears more conclusive because it enabled an overview of the spoilage phenomenon, in particular to identify similarities or differences between the two types of meat as well as to go further in understanding the underlying metabolic pathways.

When studying microbial meat spoilage, it is difficult to precisely obtain the confirmation of “who is doing what” in a complex microbial community, even if, independently, the metabolites produced at the end of storage are often well known and characterized (Casubari et al., 2015; Remenant et al., 2015). However, we were able to make very interesting observations about metabolic interactions because our holistic approach that integrated metabolomics and metataxonomics with recent statistical tools, improved the understanding of spoilage phenomenon by identifying relevant process-specific biomarkers and by correlating them with specific metabolites and spoilage phenotypes. Reconstruction of metabolic ecological networks associated with bacterial species and environmental parameters also revealed the possible modulation of burden sharing within the community members.

To start with, we revealed the possible connection of *Vibrionaceae* in isopropanol production from acetone, a pathway which may involve a secondary alcohol dehydrogenase (SADH; EC:1.1.1.80; K23259), an enzyme that is encoded in many genomes including those of *Vibrionaceae,* such as species of *Photobacterium* and *Vibrio*. According to our data (Figure 8), acetone is naturally present in raw meat batter. Secondly, *L. fuchuensis* and *L. piscium* exhibited metabolic activities associated with sulfur compound production such as methanethiol and dimethyl sulfide, respectively. Such activities are often associated with species members of *Enterobacteriales* like *Serratia* or *Hafnia* (Casubari et al., 2015). We partly confirmed this view, whereas *L. fuchuensis* and *L. piscium* show the strongest positive correlation with the production of these spoilage-related metabolites. Although *L. piscium* spoilage of pork meat has been reported as characterized by buttery and sour odors (Remenant et al., 2015,)), our observation is interesting because it echoes previous data on the main role of another *Lactococcus* species, *Lactococcus lactis,* and its role in sulfur compound production from methionine or cysteine in cheese (Hannify et al., 2009). Furthermore, within the genus *Latilactobacillus*, our data show that *L. fuchuensis* would represent a species with greater spoilage potential than *L. sakei* and *L. curvatus* (Remenant et al., 2015) due to the production of these sulfur compounds from methionine catabolism.

The construction of an ecological network also offers the advantage of being able to connect all the metabolic pathways and the different actors in an overall view. For instance, it is possible to observe the central position of *Leuconostoc* species in the network (see Figure 10) and, in particular, the link that these species play between aerobiosis- and anaerobiosis-induced catabolic pathways. In this way, we have been able to cross-reference several observations that lead us to propose the potential existence of a commensal link between two of the most frequent species for both sausages: *L. carnosum* (with possibly *L. gelidum*) and *L. curvatus*. This hypothesis links two different pathways: the acetoin production pathways and the propanediol dehydrogenase pathway (PDU pathway).

Acetoin production follows two different pathways in bacteria, one taking place under aerobiosis conditions through spontaneous decarboxylation of acetolactate into diacetyl, followed by conversion to acetoin (Papadimitriou et al., 2016). Our results show that *B. thermosphacta, Pseudomonas* species*, Psychrobacter cibarius* and *Leuconostoc* species are the main producers of acetoin during aerobiosis in raw sausages, and these data corroborate many previous studies on meat spoilage (Stanley et al., 1981; Casubari et al., 2015, Illikoud et al., 2019; Papadopoulou et al., 2020). The second pathway is instead an anaerobic process that takes place under low carbohydrate availability through acetolactate decarboxylation by the ALD (α-acetolactate decarboxylase) enzyme to acetoin and conversion of acetoin to 2,3-butanediol by the BDH (butanediol dehydrogenase) enzyme (Papadimitriou et al., 2016). In this pathway, 2,3-butanediol is therefore the end product because its production provides additional NADH reoxidation to the bacterial cells. *Leuconostoc* species are known to use this pathway quite actively, in particular, during citrolactic fermentation (Cogan and Jordan, 1994; Marty-Teysset et al., 1996; Zaunmuller et al., 2006). Interestingly, 2,3 butanediol could not be identified in the sausage samples as VOC using GC-MS analysis, whereas such a molecule should have been detectable with our metabolomic methodology. On the other hand, *Leuconostoc* species were positively correlated with 2-butanone and 2-butanol production under anaerobic conditions. Using these observations, we made the link with the PDU pathway.

The propanediol dehydrogenase or PDU pathway is encoded by a large *pdu* gene cluster in bacteria. This cluster is responsible for the production of the enzymatic complex and its cognate intracellular polyhedral body formation (Bobik et al., 1999). The pathway converts 1,2 propanediol to 1-propanol or to propionate. The substrate, 1,2 propanediol, may have several origins in bacteria but the main one is from glycerol and DHAP catabolism via the methylglyoxal reductase pathway and is often associated with phospholipid catabolism. Furthermore, a recent study showed that the PDU pathway displays broad substrate specificity in *Lactobacillaceae* and that the large protein complex is capable of converting 2,3 butanediol into 2-butanol (Russmayer et al., 2019). A detailed mining of the genomes of the many strains from the various species identified in our sausage bacterial communities (data not shown) revealed that only strains from *L. curvatus* may possess the *pdu* gene cluster (Terán et al., 2018), which is then absent from the *L. carnosum* and *L. gelidum* genomes that we analyzed. Finally, the ecological network shown in Figure 10 clearly establishes a positive correlation between *L. curvatus* with both 1-propanol and 2-butanol production.

The most straightforward explanation arising from these observations is that 2,3 butanediol is the molecule that links both pathways and that provides the possible commensal relationship between *L. carnosum/gelidum* and *L. curvatus*. Under anaerobiosis, *Leuconostoc* species would convert acetoin to 2,3 butanediol, which is then used as a substrate by *L. curvatus* to produce 2-butanol. Both members then receive the mutual beneficial NADH reoxidation in the process. From our point of view, this hypothesis also provides an explanation of the very frequent domination of raw sausage microbial communities by these two species.

However, these assumptions made thanks to statistical correlations between heterogeneous data still need to be assessed. Nevertheless, our observations open great perspectives for synthetic ecology approaches (for example, the reconstruction of microbial consortia to decipher metabolic interactions) in order to reproduce and decipher this trophic relationship. The use of metagenomics and metatranscriptomics would also be of great interest to characterize the response of these communities to variations in environmental and process parameters.

Our study also highlights the limitations associated with single gene-based amplicon sequencing (either 16S-based or *gyrB*-based as in the present study) to describe spoilage phenomena. To enrich our knowledge of this problem and provide predictive tools for reliable estimation of the shelf life of fresh meat products, a holistic approach that combines the description of detailed structuration and activity (gene expression) of microbiota in various environmental conditions is needed. Industrials must be involved in this process to foster large sampling campaigns in representative contamination conditions.

---

**Supplementary Figure S1.**
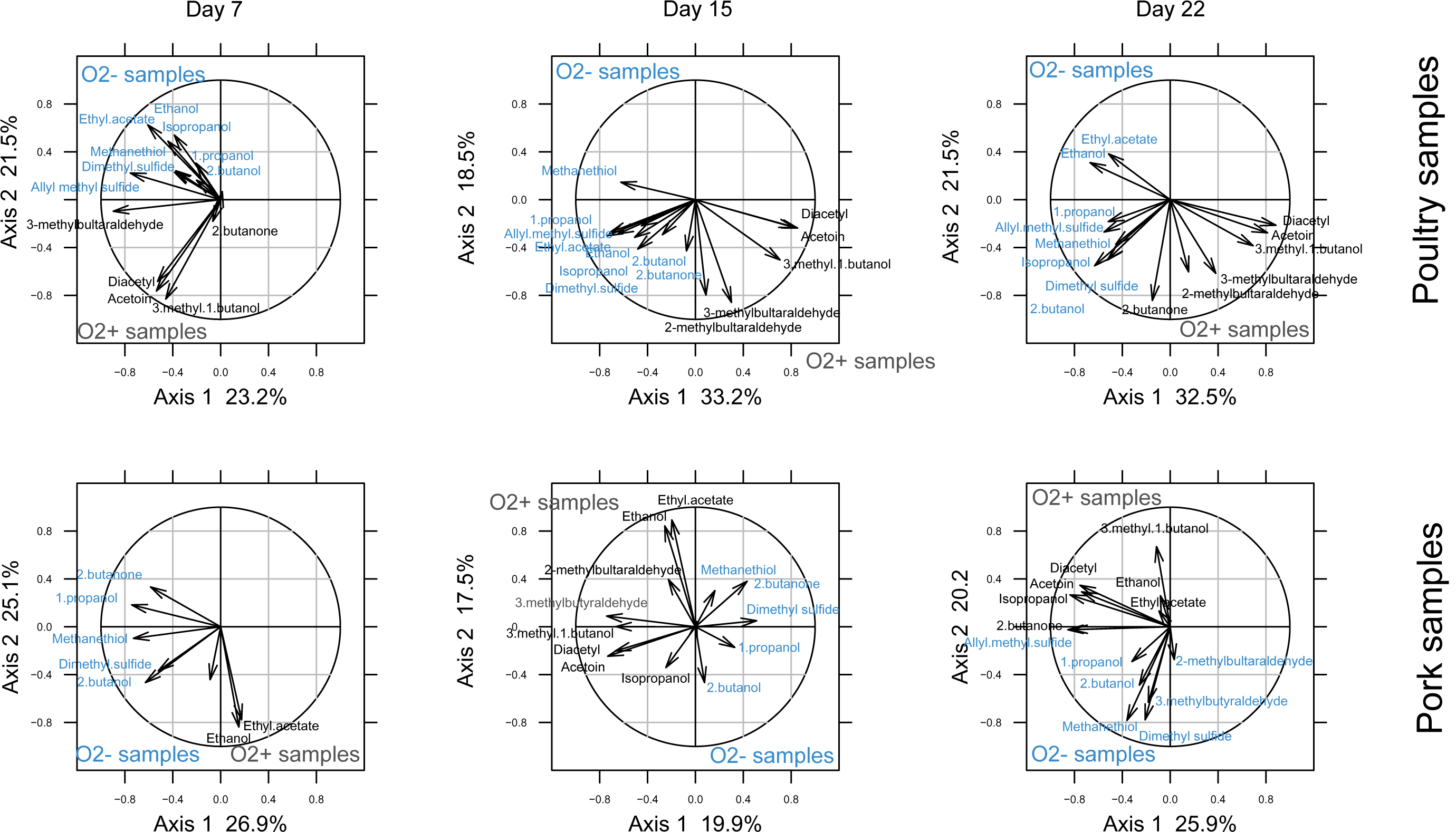
Correlation circles highlighting the role of the VOCs in the distribution of the samples in the factorial plane shown in Figure 9. Poultry samples are shown on the top and pork samples at the bottom. From left to right, correlation circles correspond to storage time from 7 to 22 days. Colors for VOC descriptors correspond to O_2_+ (dark gray) and O_2_-(blue) storage conditions.

**Supplementary Figure S2.**
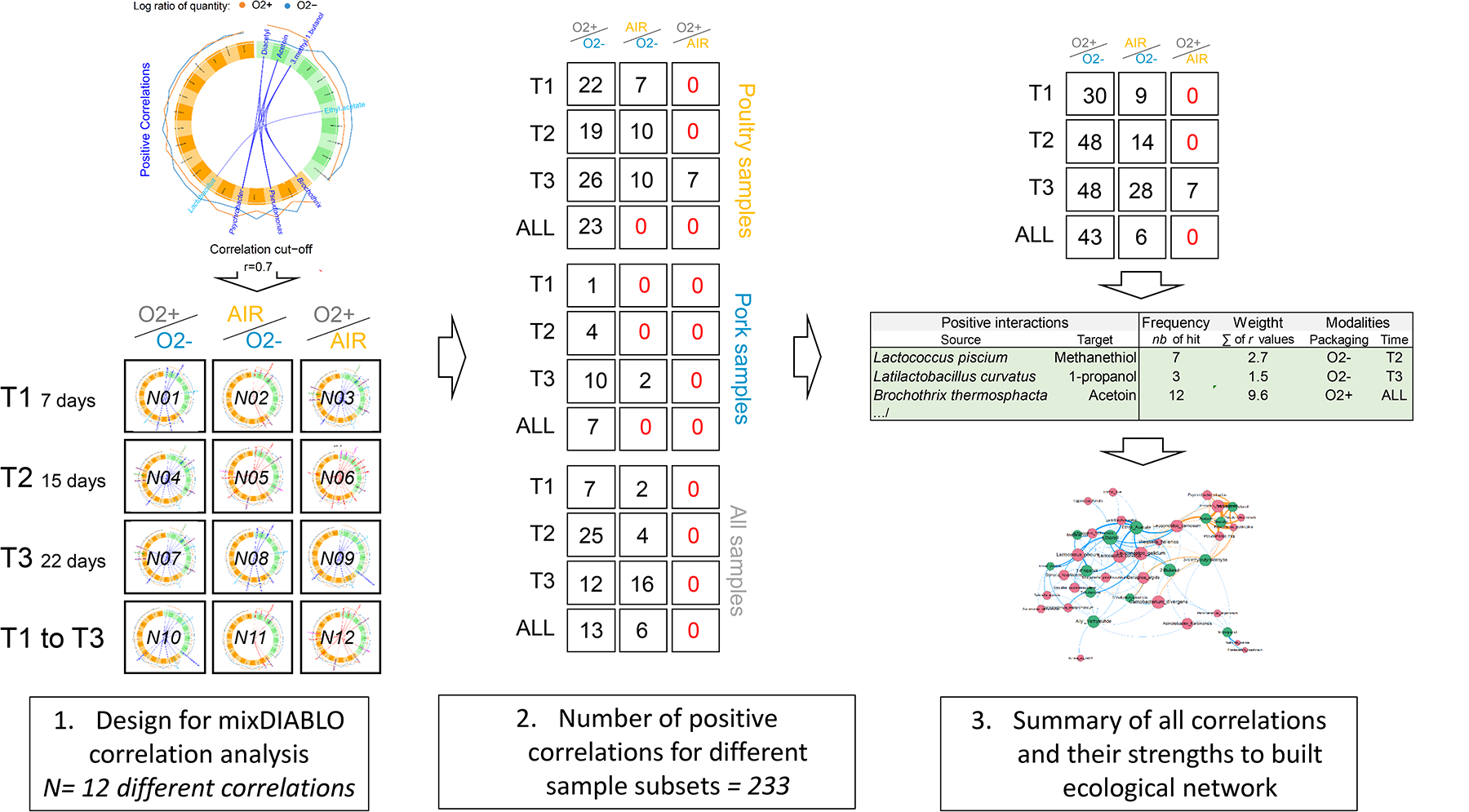
Design and results of the variable-driven subset mixDIABLO analyses. The first step (left upper part) illustrates how positive correlation (blue lines) between bacterial species (yellow) and metabolites (green) were selected between two packaging conditions. In this figure, for example only, the r value cut-off was set at 0.7 (but ranged from 0.5 to higher values) and taxons are presented at the genus level. In a similar way (left bottom part), analysis of 12 different subsets could be designed depending on the packaging comparison (columns) and the time of storage (rows) including subsets where all storage times were combined. This design was applied to the dataset from the two meat types (pork or poultry) and when both meats were combined (central part). This resulted in the analysis of 36 different subsets, yielding, overall, 233 positive correlations between bacterial species and metabolites. All correlations were summarized into a table for ecological network construction (right part). Some positive correlations were identified in several subset analyses, whereas some others were found in only a few of them. For instance, positive correlation between *B. thermosphacta* and acetoin was found in 12 subsets (out of 36) with cumulated r values of 9.6 (average r value of 0.8 per positive correlation). Similarly, positive correlation of *L. curvatus* with 1-propanol was found in three subsets (only at T3 of storage) with an average r value of 0.5 per positive correlation.

**Supplementary Figure S3.**
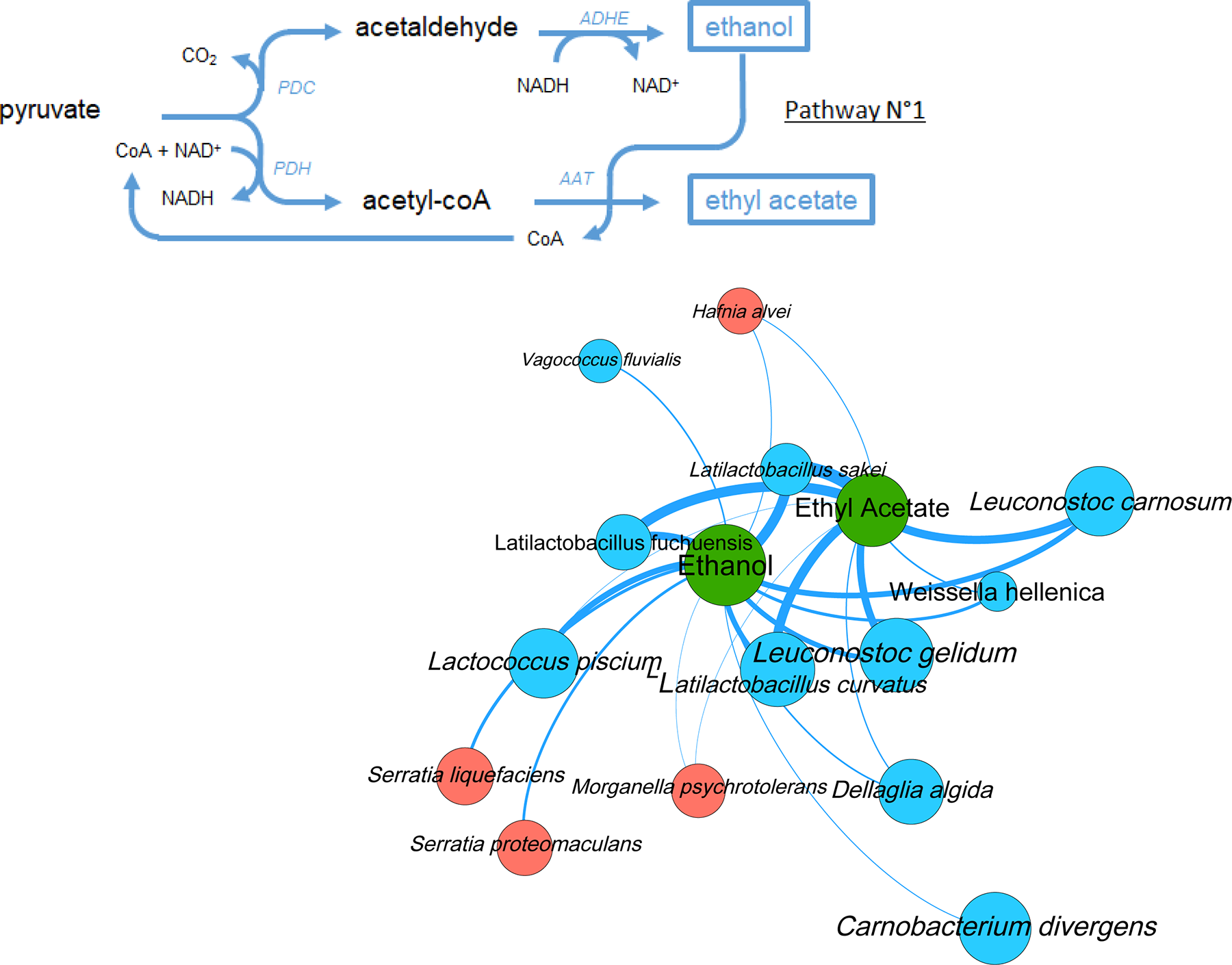
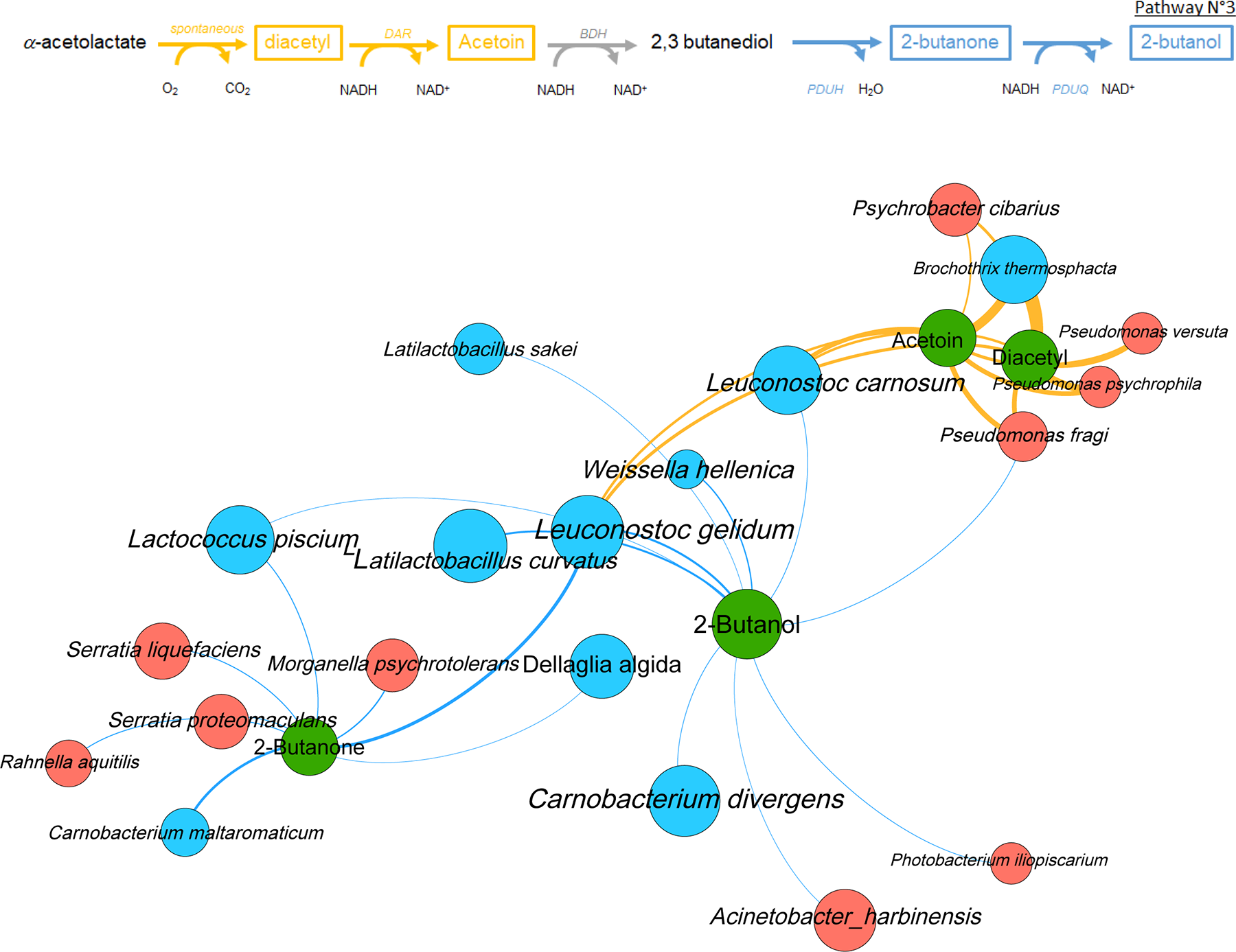
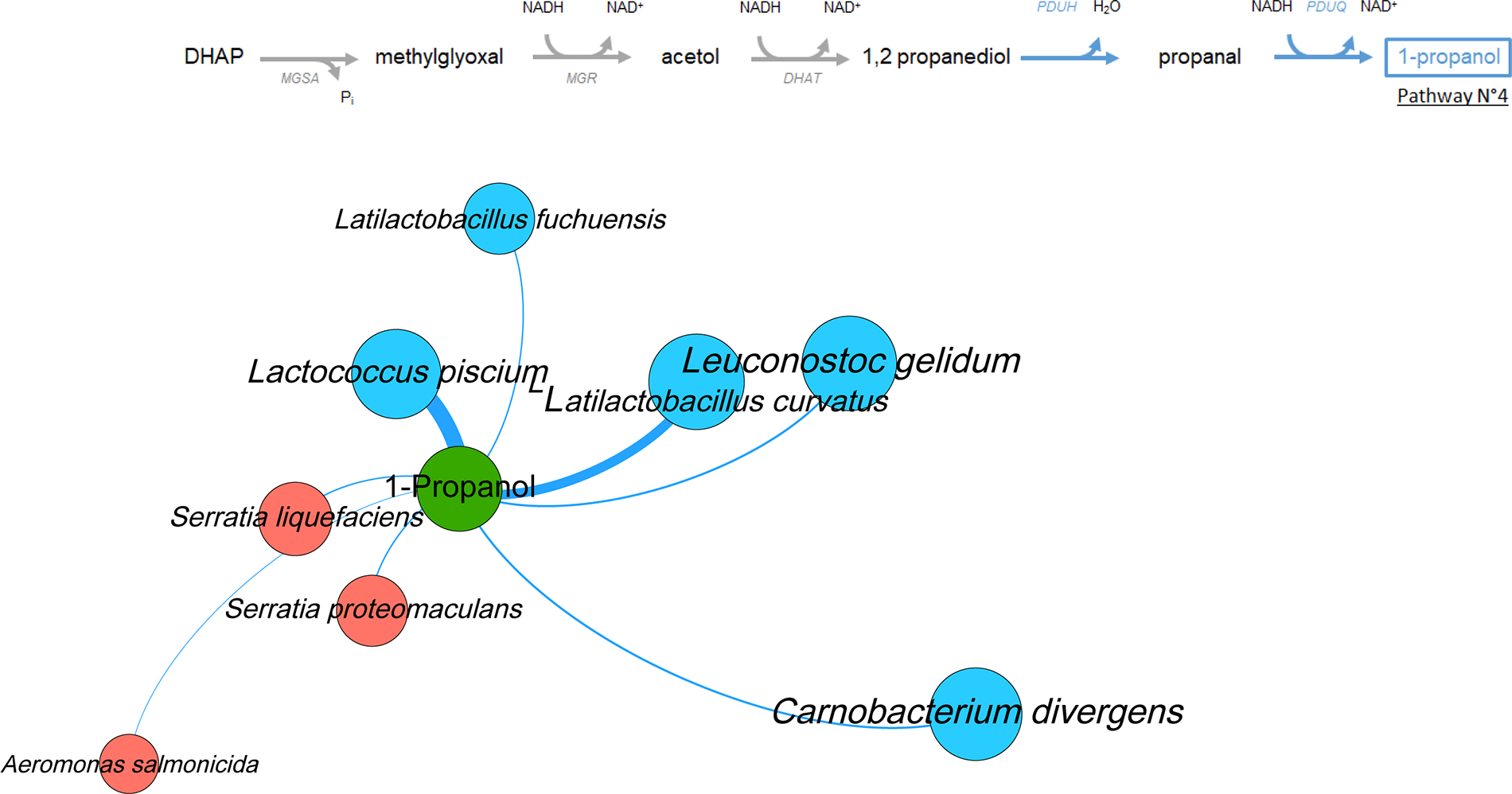
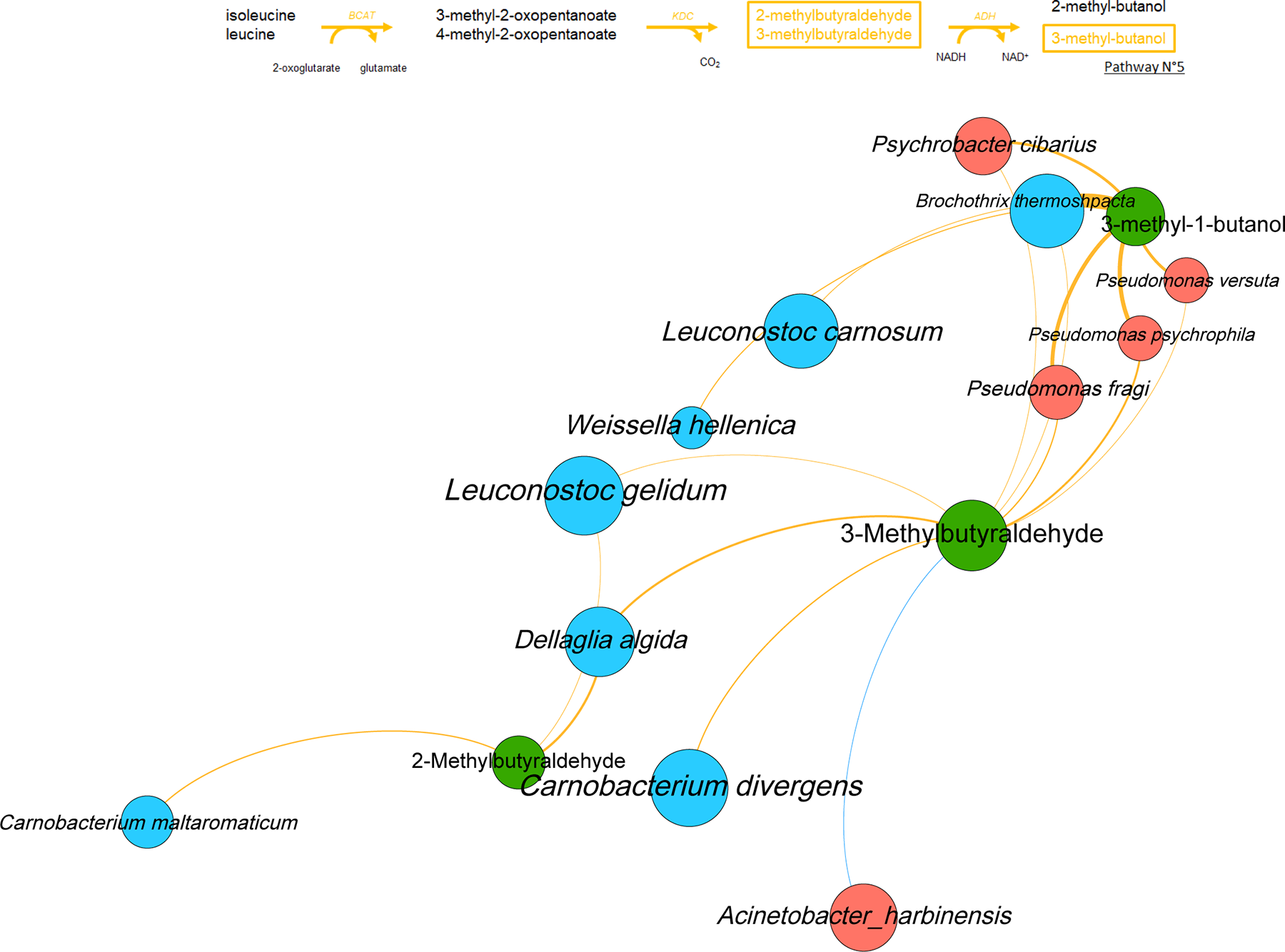
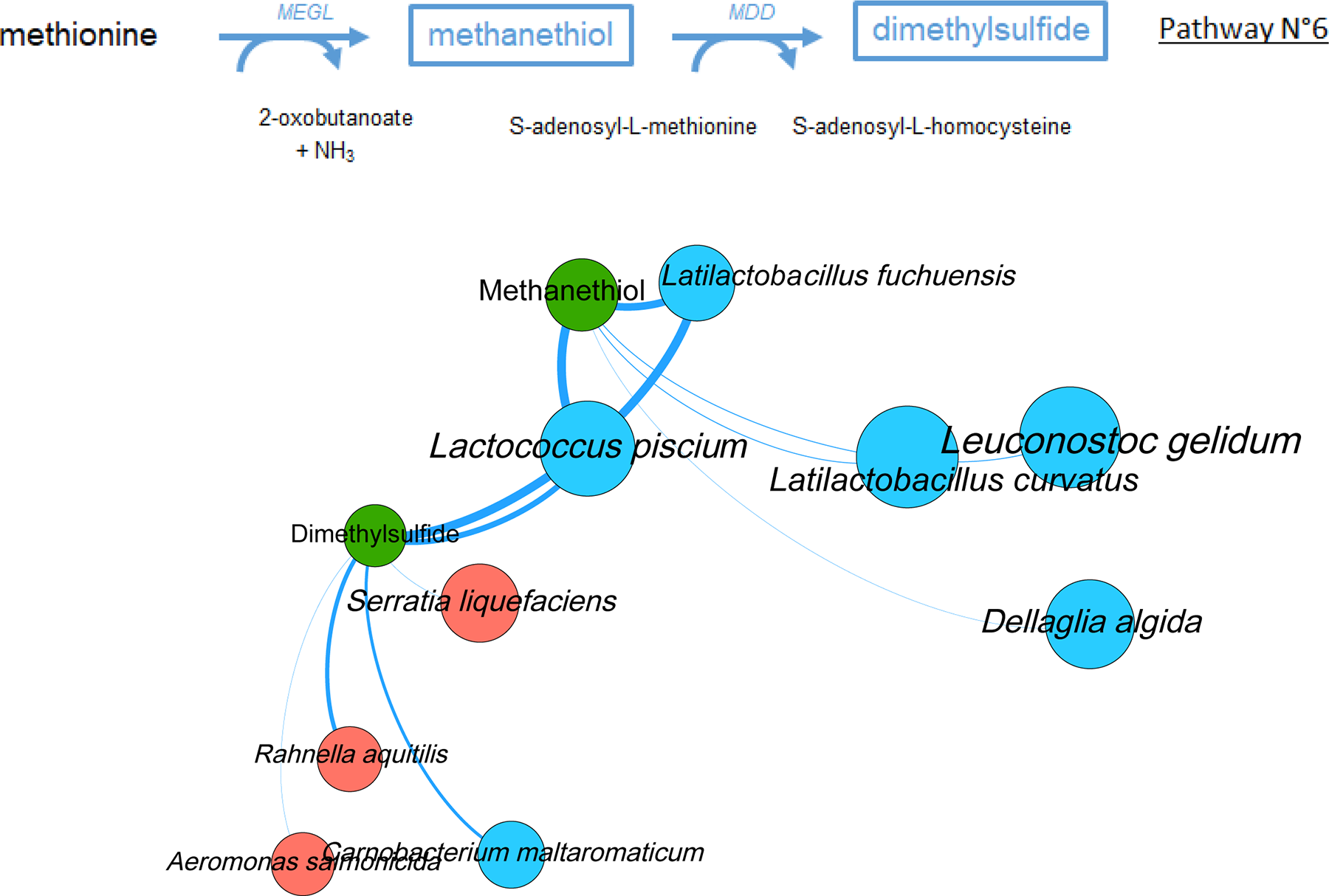
This supplementary figure is a series of extractions from Figure 10 independently showing the six metabolic pathways and the ecological network that is associated with them. Legends are as in Figure 10.

## Acknowledgments

This work was supported by the ANR Redlosses Project, Grant ANR-16-CE21-0006, overseen by the French National Research Agency (ANR). SP was granted a postdoctoral through this agency. The funding body did not play any role, neither in the design of the study nor in the collection, analysis or interpretation of data. The authors express their gratitude to the INRAE MIGALE bioinformatics facility *(MIGALE, INRAE, 2022. Migale bioinformatics Facility,* doi: 10.15454/1.5572390655343293E12) for providing computational resources and data storage.

## ANR Relosses consortium group members

Simon Poirier, Ngoc-Du Martin Luong, Valérie Anthoine, Sandrine Guillou, Jeanne-Marie Membré, Nicolas Moriceau, Sandrine Rezé, Monique Zagorec, Carole Feurer, Bastien Frémaux, Sabine Jeuge, Emeline Robieu, Marie-Christine Champomier-Vergès, Gwendoline Coeuret, Emilie Cauchie, Georges Daube, Nicolas Korsak, Louis Coroller, Noémie Desriac, Marie-Hélène Desmonts, Rodérick Gohier, Dalal Werner, Valentin Loux, Olivier Rué, Marie-Hélène Dohollou, Tatiana Defosse, Stéphane Chaillou.

## Notes

https://doi.org/10.15454/UDQLGE

### Competing Interest Statement

The authors have declared no competing interest.

